# CNVision: a computational framework for functional interpretation of intragenic copy number variants

**DOI:** 10.64898/2026.10.28.762641

**Authors:** Miles Liao, Xinming Zhuo

## Abstract

**Background:** Intragenic copy number variants (CNVs), including exon-level deletions and duplications within a single gene, represent an important class of pathogenic genomic variation. Their functional consequences depend on the precise relationship between the CNV breakpoints, exon structure, coding sequence boundaries, transcript orientation, and reading frame. Although intragenic CNVs contribute substantially to Mendelian disease, their interpretation remains difficult because many existing workflows rely on manual transcript review or annotation pipelines that do not consistently resolve exon-level and base-pair-level consequences.

**Results:** We developed CNVision, a containerized computational framework for standardized interpretation of intragenic CNVs. CNVision accepts multiple input formats, including exon-based entries, genomic coordinates, and HGVS genomic notation, and converts each into a unified internal representation for transcript-aware analysis. The system maps CNVs to coding sequence regions, calculates affected coding sequence length at base-pair resolution, and classifies predicted reading-frame consequence. CNVision was evaluated using a dataset including 200 curated intragenic CNVs spanning multiple genes, genome builds, variant structures, and input types. Across the dataset, CNVision produced predictions concordant with manual transcript-level analysis and correctly processed exon-based, coordinate-based, and HGVS-formatted inputs. Beyond basic evaluation, CNVision also identified discrepancies in previously published CNVs. Of 131 reported CNVs evaluated, 15 yielded classifications that differed from the original published interpretation; independent manual transcript-level analysis using the UCSC Genome Browser supported CNVision’s classification in all 15 variants, indicating the discrepancies were attributable to differences in transcript interpretation or coordinate calculation rather than computational error. In terms of processing speed, CNVision was tested by two board-certified laboratory geneticists who compared the runtime of CNVision versus manual analysis across 18 CNVs, 6 of each input type. CNVision consistently outperformed manual analysis by both geneticists across all variants by a significant margin.

**Conclusion:** CNVision provides a reproducible framework for functional interpretation of intragenic CNVs by automating input parsing, transcript mapping, coding sequence overlap calculation, and reading-frame classification. The framework reduces manual burden, improves consistency, and provides a foundation for future expansion into batch CNV processing and graphical variant interpretation.

## Background

Intragenic copy number variants (CNVs), also referred to as exonic or monogenic CNVs, are a significant source of genomic variation defined as deletions or duplications of genetic material within a single gene. These alterations vary widely in scope, ranging from sub-kilobase changes involving only a single exon to large-scale events affecting multiple exons or the entire gene [1,2].

The molecular and functional consequences of intragenic CNVs are diverse and depend heavily on the nature and location of the alteration. Exonic deletions and duplications often result in a shift in the reading frame, introducing premature stop codons that trigger nonsense-mediated decay (NMD), effectively creating a null or loss-of-function allele. In other instances, these variants may escape NMD, resulting in truncated proteins with dominant-negative effects. Conversely, in-frame losses or gains can alter protein length and structure, potentially leading to hypomorphic or even neomorphic protein functions. Beyond the coding sequence, intragenic CNVs can involve regulatory elements such as promoters, mimicking whole-gene deletions by impairing gene expression and function.

The identification of intragenic CNVs has surged due to the expanding use of high-resolution microarrays and next-generation sequencing (NGS). Transitioning from low-resolution karyotyping and fluorescence in situ hybridization (FISH), clinical laboratories adopted targeted, exon-level array comparative genomic hybridization (aCGH) platforms capable of detecting sub-kilobase copy number changes [1,3,4]. More recently, the widespread adoption of whole-exome sequencing (WES) and whole-genome sequencing (WGS) has enabled the simultaneous detection of single-nucleotide variants (SNVs) and CNVs within a single assay, representing a paradigm shift toward comprehensive genomic testing [2,5–8].

Recent large-scale clinical studies further highlight the importance of intragenic CNVs in molecular diagnosis. Intragenic CNVs were identified in 9.8% of individuals receiving a clinical report containing a pathogenic or likely pathogenic variant and accounted for 4.7–35% of pathogenic variants depending on clinical specialty [9]. In an exon-targeted microarray study of 18,708 patients, 8,094 (33.2%) of 24,373 non-polymorphic CNVs involved only a single gene [10]. Similarly, in a pediatric cohort of 1,032 individuals undergoing clinical genome sequencing, 32.9% of reported deletions overlapped only a single gene [7].

Despite their established clinical significance, the interpretation of intragenic CNVs remains challenging. Existing annotation pipelines often lack the transcript-aware precision required for accurate interpretation, particularly for variants involving only one or a few exons. To address these limitations and provide a scalable framework for standardized intragenic CNV analysis, we developed CNVision. CNVision integrates standardized genomic resources with an automated computational workflow to accurately assess coding sequence disruption, predict reading-frame consequence, and support consistent functional interpretation of intragenic CNVs.

## Implementation

### System architecture

CNVision was implemented as a modular, web-accessible computational framework for intragenic CNV interpretation. The system consists of a browser-based user interface, a Flask-based backend server, and a Python analysis pipeline responsible for input parsing, transcript mapping, coding sequence analysis, and result generation. The interface was developed using HTML, CSS, and Bootstrap to provide a lightweight and accessible entry point for users, while the backend was written primarily in Python to support flexible genomic data processing and rapid development of parsing logic.

The application follows a client-server design. Users submit CNV information through the web interface, where the selected input mode determines the required fields and expected notation structure. The Flask backend receives the submitted form data, validates the input, routes the request to the appropriate parser, and passes the normalized variant representation into the downstream analysis pipeline. The major computational stages include input parsing, input normalization, gene and transcript resolution, exon and coding sequence mapping, frame consequence classification, and structured result display (Figure 1).

**Figure 1.**
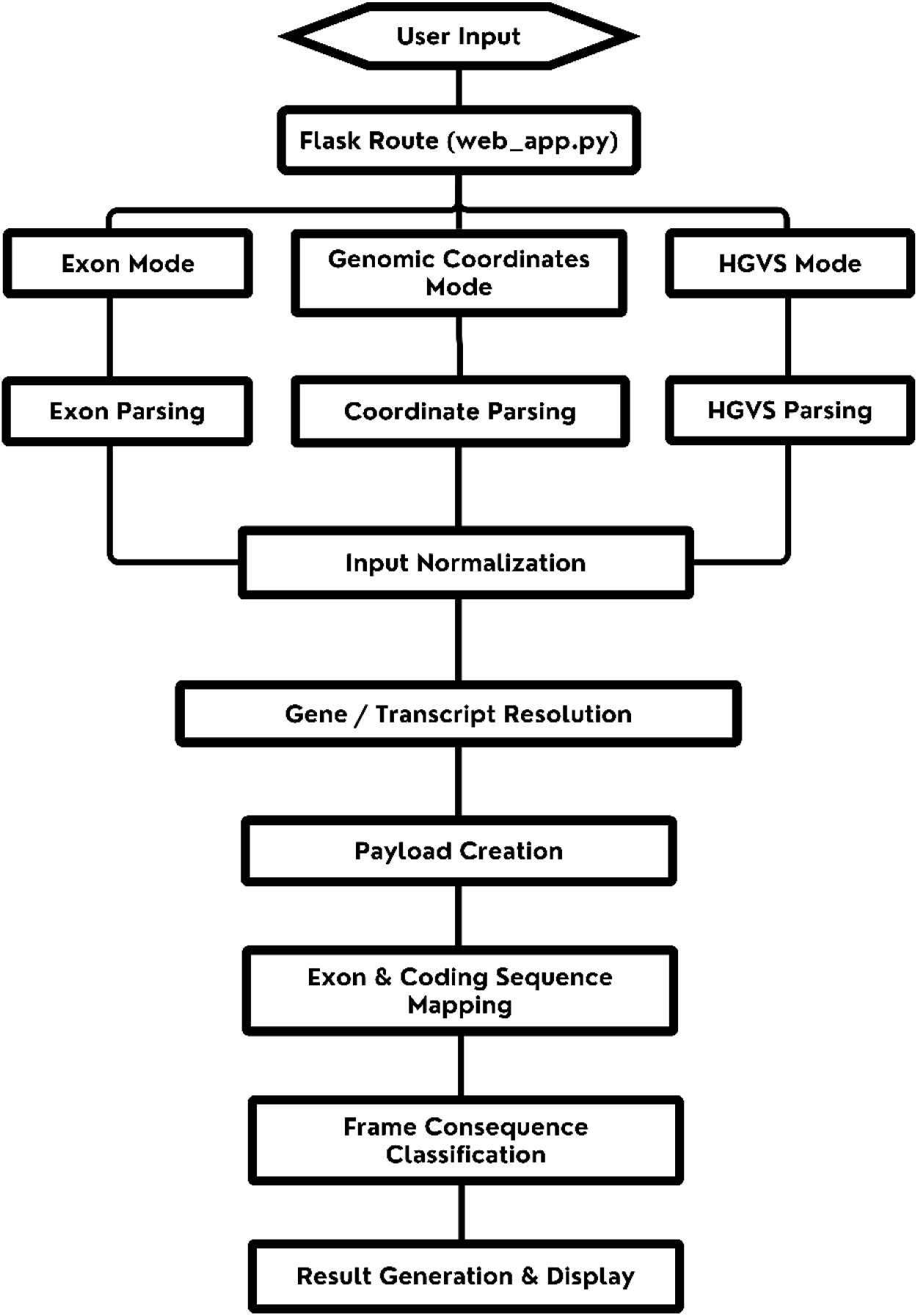
Overview of the CNVision computational workflow. User-submitted intragenic CNVs are parsed from supported input formats, validated and normalized, mapped to transcript and CDS annotations, analyzed for coding sequence overlap, classified according to predicted reading-frame consequence, and returned through the web interface.

To support reproducibility and deployment across environments, CNVision was containerized using Docker and deployed through Microsoft Azure App Service. Containerization ensures that the application, software dependencies, runtime configuration, and analysis environment remain consistent across development and production environments, while Azure App Service provides a managed hosting platform for reliable web-based access.

CNVision was developed using Python 3.10.18 with Flask 3.1.1 as the web framework. The user interface was implemented using HTML5, CSS3, and Bootstrap 5.3.7. The application was containerized using Docker 28.3.0 to ensure reproducible deployment across computing environments.

### Input modes and variant representation

CNVision supports three primary input modes: exon-based input, genomic coordinate/ISCN nomenclature [11] input, and HGVS nomenclature [12] input. These formats were selected because they represent the most common ways intragenic CNVs are reported in research studies, clinical laboratories, variant databases, and microarray analyses. Regardless of the input format, all variants are converted into a standardized internal representation before downstream analysis.

### Exon-based input

In exon-based mode, the user provides a gene symbol and the affected exon or exon range. CNVision automatically selects the corresponding MANE Select transcript [13] whenever one is available; otherwise, the RefSeq Select transcript [14] is used. The submitted exon numbers are then mapped to the selected transcript’s exon and coding sequence (CDS) coordinates. This mode is intended for cases in which CNVs are reported at the exon level, such as "deletion of exon 5" or "duplication of exons 3–7," without precise genomic breakpoint information.

### Genomic coordinate input

In genomic coordinate mode, the user provides a chromosome number, start coordinate, end coordinate, genome build, optional gene symbol, and CNV type. CNVision validates the submitted coordinates and maps the genomic interval directly to the selected transcript to determine the affected coding sequence. CNVision also accepts microarray results directly in the ISCN nomenclature. For example, arr[GRCh37] 7q11.22(70204489_70236787)x3. Because this mode parses specific coordinates, it can fully handle partial exon deletion or duplication as well. This mode is designed for variants identified by sequencing, chromosomal microarray, or genomic databases where genomic breakpoints are available.

### HGVS nomenclature input

CNVision accepts genomic variants described using HGVS nomenclature. Users can directly paste standardized HGVS variant descriptions into the application without manual reformatting. For example, NC_000015.9:g.(42652323_42676657)_(42684925_42686351)del. Because this mode parses specific coordinates, it can fully handle partial exon deletion or duplication as well. CNVision extracts the relevant genomic coordinates and variant information from the submitted notation, infers the appropriate reference information when possible, and converts the variant into the standardized internal representation used for downstream analysis.

### Variant normalization

After parsing, all input types are converted into a unified internal representation containing the selected transcript, gene symbol, variant type, genome build, genomic interval (when available), affected exon range (when applicable), and parsed notation metadata. This standardized representation enables the same transcript mapping and consequence prediction workflow to be applied consistently regardless of the original input format.

### Reference transcript selection and annotation resources

CNVision performs transcript-aware analysis using curated transcript annotation resources. The primary transcript reference is based on MANE transcripts when available, with RefSeq Select transcripts used as a secondary reference where appropriate. Transcript records include gene symbols, transcript identifiers, chromosome, strand orientation, exon boundaries, coding sequence start and end positions, and exon ordering.

Transcript selection is a critical step because the predicted consequence of an intragenic CNV can depend on the transcript model used for interpretation. CNVision, therefore, uses a standardized transcript selection strategy rather than relying on arbitrary user selection. For each submitted gene, the framework resolves the corresponding reference transcript and retrieves the exon and coding sequence structure required for downstream analysis. If the gene cannot be resolved or if required transcript metadata are unavailable, the system returns an interpretable error rather than producing an unsupported classification. For example: “Gene ‘ABC’ not found in MANE GRCh38 data”

CNVision accounts for transcript strand orientation during exon and coding sequence mapping to ensure accurate interpretation of transcript-specific genomic coordinates.

### Input normalization and quality control

Before functional analysis, CNVision performs input normalization and organization. The purpose of this stage is to ensure that all variants are represented consistently and that downstream calculations are performed only on valid inputs. The normalization process includes standardizing chromosome labels, checking coordinate types, validating genome build selection, resolving gene symbols, confirming that exon numbers exist within the selected transcript, and ensuring that the submitted interval overlaps the expected gene region when coordinate information is available.

For coordinate-based and HGVS-based inputs, CNVision checks whether the start and end positions form a valid interval and whether the interval can be evaluated against the selected transcript. For exon-based inputs, the system verifies that the submitted exon range falls within the transcript’s exon count. If a user submits an invalid exon number, malformed notation, unsupported genome build, or gene symbol that cannot be resolved, CNVision returns a structured error message rather than producing a potentially misleading prediction. For example: “Exon range 200-200 exceeds transcript bounds. NM_004006.3 has 79 exons.”

This validation process is essential because intragenic CNV interpretation depends on accurate mapping of variant coordinates to transcript and coding sequence boundaries. Errors in coordinate specification, transcript selection, or exon assignment can affect coding sequence overlap calculations and, consequently, the predicted functional consequence. CNVision therefore incorporates explicit validation and deterministic processing steps to ensure consistent and reproducible classification prior to downstream analysis.

### Exon and coding sequence mapping

After transcript resolution, CNVision maps the submitted CNV to the exon and coding sequence structure of the selected transcript. For exon-based inputs, the affected exon range is converted into genomic intervals using the transcript annotation. For coordinate-based and HGVS-based inputs, the submitted genomic interval is directly intersected with transcript exon intervals.

CNVision distinguishes between total exon overlap and coding sequence overlap. This distinction is necessary because not all exonic sequences are coding. Some exons contain untranslated regions, and some CNVs may overlap only non-coding portions of a transcript. For each affected exon, CNVision identifies the coding portion of the exon by intersecting the exon interval with the transcript’s coding sequence start and end coordinates. The variant interval is then intersected with this coding portion to calculate the number of coding base pairs affected.

The system performs this calculation at base-pair resolution for each affected exon. For complete exon events, the coding overlap corresponds to the coding portion of the full exon. For partial exon events, the coding overlap corresponds only to the intersection between the CNV interval and the coding portion of the exon. The affected coding sequence length is calculated as the sum of coding overlap across all affected exons.

### Reading-frame consequence classification

CNVision classifies predicted reading-frame consequences using the total affected coding sequence length. Once the coding sequence overlap has been calculated, the framework evaluates whether the number of affected coding base pairs is divisible by three. If the affected coding length is divisible by three, the CNV is classified as predicted in-frame. If the affected coding length is not divisible by three, the CNV is classified as a predicted frameshift. If the CNV does not overlap the coding sequence, it is classified as non-coding with respect to the selected transcript.

This modulo-based approach provides a deterministic and transparent method for predicting reading-frame disruption. The rule is directly tied to codon structure, because each codon consists of three nucleotides. A deletion or duplication that changes the coding sequence length by a multiple of three is expected to preserve the downstream reading frame, whereas a length change not divisible by three is expected to shift the reading frame downstream of the event.

CNVision reports the affected coding sequence length, the modulo result, and the final predicted consequence. This transparency allows users to verify how the classification was produced rather than relying on an opaque output label. The framework is designed to support functional interpretation, not to replace clinical pathogenicity classification [15, 16]. Therefore, outputs are best understood as predicted molecular consequences that can support downstream variant interpretation. CNVision does not predict other functional effects, such as disruption of start or stop codons, alternative translation initiation, or protein truncation mechanisms that extend beyond reading-frame analysis.

### Result generation and user interface

CNVision generates structured results that summarize the submitted variant, transcript used for analysis, affected exons, coding sequence overlap, base-pair calculations, and predicted reading-frame consequence. The result display is designed to show both the final classification and the intermediate computational steps used to generate it.

For each analyzed CNV, the output includes the input mode, gene, genome build, transcript identifier, variant type, affected genomic interval when available, affected exon or exon range, total coding sequence length affected, and predicted functional consequence. When applicable, the output also identifies partial exon overlap and displays exon-level coverage information. This allows users to distinguish between complete exon events and events that affect only part of an exon or coding region.

The user interface was designed to make CNVision usable without requiring command-line execution (Figure 2A). Users can enter CNV information through the browser, select the input type, submit the variant for analysis, and review the resulting prediction. The current interface supports single-variant analysis and displays results in a human-readable format, with representative output shown in Figure 2B. The underlying data structure is compatible with future expansion toward machine-readable outputs and batch processing.

**Figure 2.**
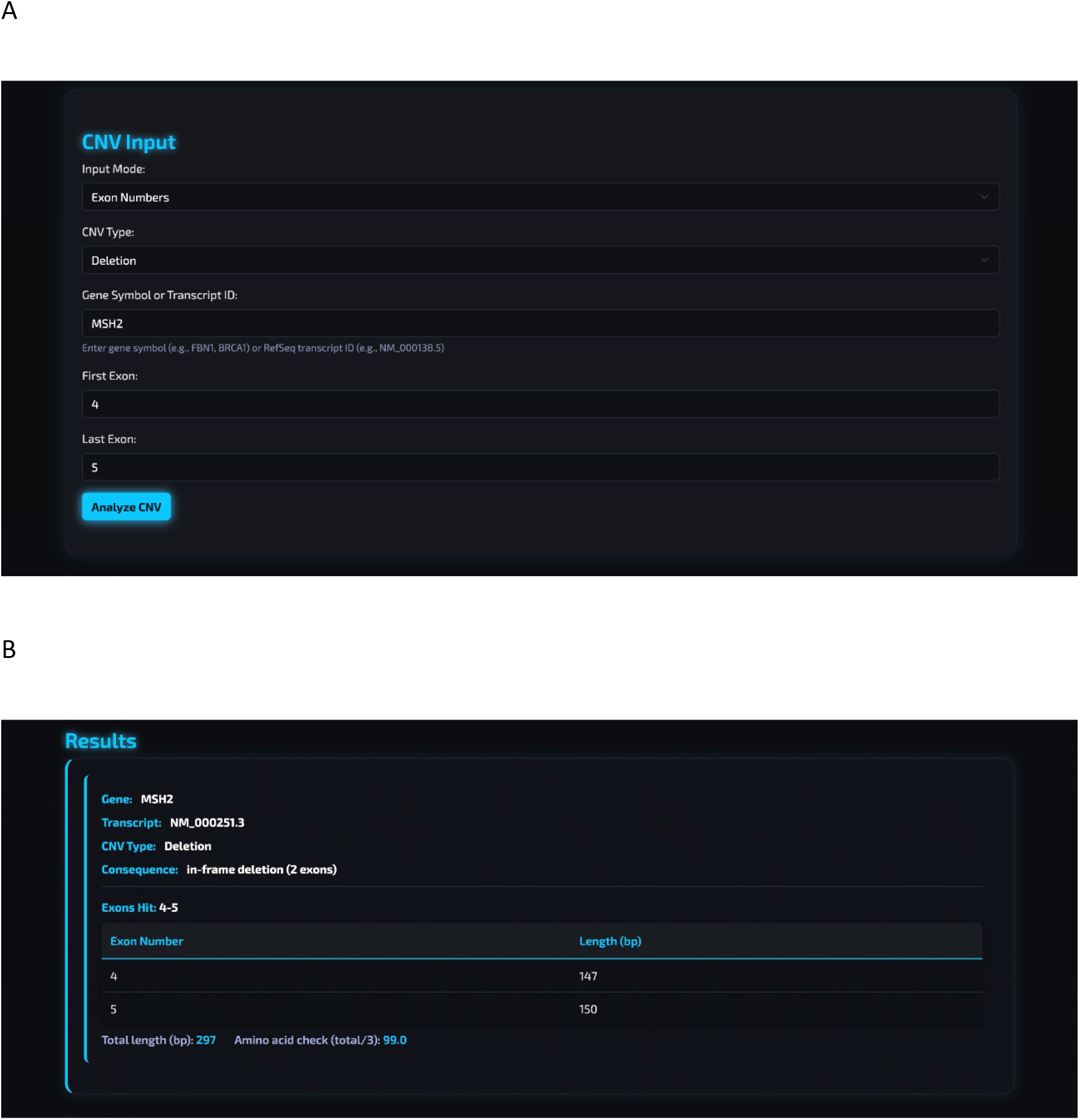
CNVision web interface. A. Representative intragenic CNV submission. The browser-based interface supports exon-based, genomic coordinate/ISCN, and HGVS nomenclature input formats. Users specify the variant information and reference genome before initiating transcript-aware consequence analysis.**B. Representative CNVision analysis output.** The output summarizes the submitted variant, selected transcript, affected exon(s), coding sequence overlap, affected coding sequence length, and predicted functional consequence. Intermediate calculations are displayed to facilitate result interpretation and manual verification.

### Software organization and deployment

The CNVision codebase is organized into separate files and modules corresponding to the main functional components of the application. The Flask application version 3.1.2 handles routing and user interaction, while supporting Python modules that perform parsing, transcript lookup, coordinate conversion, coding sequence analysis, and result formatting. Static files define the visual structure and styling of the user interface.

Docker is used to package the application and dependencies into a reproducible environment. This allows CNVision to be deployed without requiring users to manually configure Python packages, server settings, or environment-specific dependencies. The containerized design also supports future deployment in institutional research environments where reproducibility, portability, and controlled execution are important.

## Results

### Overview of CNVision workflow

CNVision was developed as an end-to-end computational framework for transcript-aware interpretation of intragenic CNVs. The framework accepts exon-based descriptions, genomic coordinate intervals, and HGVS notation, which are normalized into a common internal representation prior to transcript mapping and functional consequence prediction. The overall workflow consists of input validation, transcript resolution, CDS overlap calculation, reading-frame classification, and structured result generation (Figure 1). The web-based interface enables users to submit variants using any of the three supported input formats and receive standardized functional predictions through a unified analysis pipeline (Figure 2).

### Dataset for CNVision evaluation

To evaluate the performance of CNVision, a curated dataset of 200 intragenic CNVs was assembled from multiple sources and represented the three supported input modalities (Supplementary Table). The evaluation dataset consisted of 57 exon-based variants, 101 variants described by genomic coordinates, and 42 variants reported using HGVS genomic notation. These variants encompassed 43 unique genes, 73 in-frame results, 125 frameshift results, and 2 intronic results. Of the 200 CNVs, 131 were from published literature, while the other 69 were manually crafted with diverse coding sequences and formats.

For each variant, CNVision predictions were compared against manually derived transcript-level interpretations. Manual analysis consisted of determining transcript structure, calculating the affected coding sequence length, and classifying the predicted reading-frame consequence based on coding sequence disruption. This manual interpretation served as the reference standard for evaluating CNVision performance.

### CNVision accuracy across input types

CNVision successfully processed all 200 variants included in the evaluation dataset without processing failures or unsupported input cases. Across the complete dataset, the framework achieved 100% concordance with manual transcript-level interpretation.

Performance remained consistent across all supported input modalities. CNVision correctly classified all 57 exon-based variants (100%), all 101 coordinate-based variants (100%), and all 42 HGVS-formatted variants (100%). A summary of evaluation performance across all input types is provided in Table 1.

**Table 1:** Summary of CNVision evaluation performance across three input types.

| **Input Type** | **Number of CNV** | **Correct/Incorrect** | **Accuracy** |
|---|---|---|---|
| Exon-based | 57 | 57/0 | 100% |
| Genomic Coordinate | 101 | 101/0 | 100% |
| HGVS Nomenclature | 42 | 42/0 | 100% |

For every evaluated variant, CNVision accurately determined coding sequence overlap and predicted the corresponding functional consequence based on transcript-aware reading-frame analysis. Variants involving complete exons, partial exon overlap, coding regions, and non-coding regions were all correctly classified relative to manual transcript-level analysis.

### Evaluation of previously reported CNVs

Among 200 intragenic CNVs used for evaluation, 131 were from previously published literature to assess real-world performance. Of these, 116 variants produced functional classifications that were concordant with the originally reported interpretation. Fifteen variants yielded classifications that differed from the published annotation.

Each discordant CNVs was independently re-evaluated through manual transcript-level analysis, including verification of transcript structure, coding sequence boundaries, breakpoint placement, and reading-frame calculations. Manual re-analysis supported the classifications generated by CNVision in all 15 discordant CNVs, indicating that the discrepancies were attributable to differences in transcript interpretation or coding sequence calculation rather than computational error (Table 2).

**Table 2:** Evaluation of previously published intragenic CNVs.

| Input Mode | CNV | Gene | Published Exons | CNVision Exons | Actual Exons | Published | CNVision | Actual Conse |
|---|---|---|---|---|---|---|---|---|
| Exon-based | NM_002222.6: 1-37 | ITPR1 | 1-37 | 1-37 | 1-37 | In-Frame | Frameshi | Framesh |
| Genomic Coordinates | seq[GRCh37] 19p13.2p13.2(13445205_13446729)x1 | CACNA1A | 7-8 | 7-8 | 7-8 | Frameshift | In-Frame | In-Frame |
| Genomic Coordinates | seq[GRCh37] 19p13.2p13.2(13470613_13476283)x1 | CACNA1A | 5-5 | 5-6 | 5-6 | In-Frame | Frameshift | Frameshift |
| Genomic Coordinates | seq[GRCh37] 3p26.1p26.1(4558177_4752241)x1 | ITPR1 | 1-37 | 3-38 | 3-38 | In-Frame | Frameshift | Frameshift |
| Genomic Coordinates | seq[GRCh37] 6p25.1p25.1(5368795_5369464)x1 | FARS2 | 1-2 | 2-2 | 2-2 | In-Frame | In-Frame | In-Frame |
| Genomic Coordinates | seq[GRCh37] 14q31.3q31.3(88401222_88417002)x1 | GALC | 11-17 | 12-17 | 12-17 | In-Frame | Frameshift | Frameshift |
| Genomic Coordinates | seq[GRCh37] 15q21.1q21.1(44887491_44889123)x1 | SPG11 | 24-26 | 24-26 | 24-26 | Frameshift | In-Frame | In-Frame |
| Genomic Coordinates | seq[GRCh37] 6q25.2q25.2(152685922_152690810)x0 | SYNE1 | 61-64 | 60-63 | 60-63 | In-Frame | Frameshift | Frameshift |
| Genomic Coordinates | seq[GRCh37] 2q23.3q23.3(152466318_152582063)x1 | NEB | 6-81 | 6-81 | 6-81 | In-Frame | Frameshift | Frameshift |
| HGVS | NC_000019.9:g.(13470613_13470613)_(13476283_13482502)del | CACNA1A | 5-5 | 5-6 | 5-6 | In-Frame | Frameshift | Frameshift |
| HGVS | NC_000003.11:g.(4508999_4558177)_(4752241_4753421)del | ITPR1 | 1-37 | 3-38 | 3-38 | In-Frame | Frameshift | Frameshift |
| HGVS | NC_000014.8:g.(86087482_88401222)_(88417002_88429855)del | GALC | 11-17 | 12-17 | 12-17 | In-Frame | Frameshift | Frameshift |
| HGVS | NC_000015.9:g.(44884638_44887491)_(44889123_44890374)del | SPG11 | 24-26 | 24-26 | 24-26 | Frameshift | In-Frame | In-Frame |
| HGVS | NC_000006.11:g.(152685045_152685922)_(152690810_152694177)del | SYNE1 | 61-64 | 60-63 | 60-63 | In-Frame | Frameshift | Frameshift |
| HGVS | NC_000002.11:g.(152432808_152466318)_(152582063_152584191)del | NEB | 6-81 | 6-81 | 6-81 | In-Frame | Frameshift | Frameshift |

### Runtime and computational performance

CNVision was designed for rapid single-variant interpretation through an automated transcript-aware workflow. Individual analyses were completed within timeframes under standard computational conditions. Comparison with conventional manual transcript-level interpretation by the UCSC Genome Browser [17] demonstrated a substantial reduction in analysis time while maintaining complete concordance with manual classification. Two board-certified laboratory geneticists independently evaluated the speed of CNVision versus manual analysis, as shown in Figure 3. Each geneticist was given a set of 18 cases with 6 of each input type. CNVision consistently outperformed manual analysis among 18 cases of each input mode. For laboratory geneticist #1, the range of times for manual analysis was between 22 and 99 seconds, while for CNVision it was between 10 and 16 seconds. For laboratory geneticist #2, the manual range was between 22 and 189 seconds, while for CNVision it was between 7 and 19 seconds. In terms of improvement, the average factor of improvement for geneticist #1 was 3.9x faster, while geneticist #2 showed an even more significant average improvement of 6.9x the speed of manual analysis. Case by case, the most significant improvement for geneticist #1 was a 9.9x improvement, while for geneticist #2 it was an even more impressive 27x improvement. In terms of runtime, CNVision demonstrates significant improvements compared to manual analysis using the UCSC Genome Browser.

**Figure 3.**
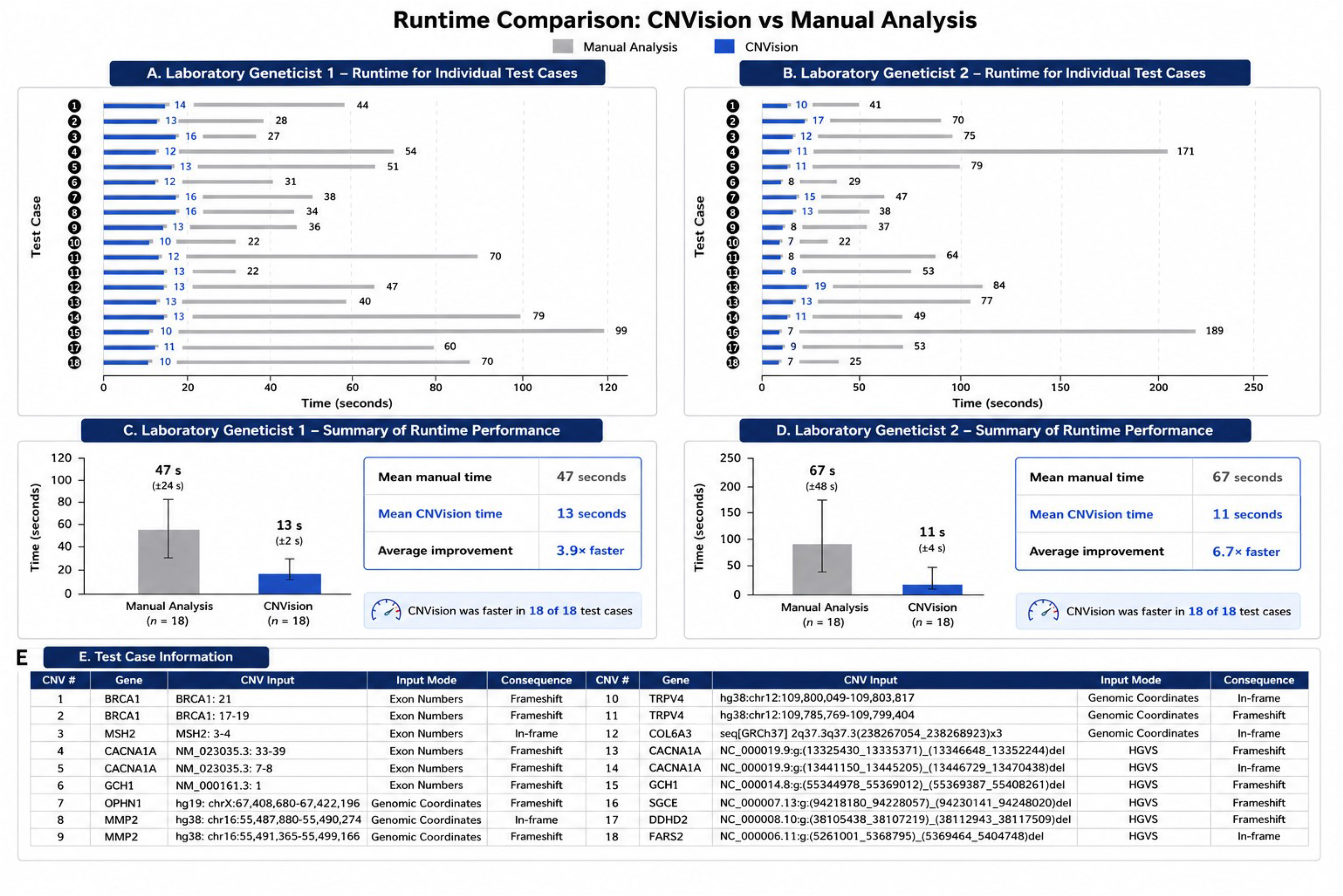
Runtime comparison between CNVision and manual transcript-level analysis. Individual analysis times for 18 test CNVs are shown for two board-certified laboratory geneticists (A, B), with mean runtimes and average speedup summarized for each geneticist (C, D). Test CNV details, including gene, input format, and predicted consequence, are listed in E.

The lightweight Python- and Flask-based architecture maintained consistent performance across exon-based, coordinate-based, and HGVS-formatted inputs without observable differences in processing time. Because the current implementation is designed for single-gene, single-variant analysis, additional benchmarking will be required following the future implementation of batch-processing functionality.

## Discussion

### Principal findings

CNVision provides a standardized computational framework for predicting the functional consequence of intragenic CNVs. The framework accepts heterogeneous input formats, resolves variants against transcript annotations, calculates affected coding sequence length at base-pair resolution, and classifies predicted reading-frame consequences using transparent modulo-3 logic. In a curated evaluation set of 200 intragenic CNVs, CNVision produced classifications concordant with manual transcript-level analysis across exon-based, genomic coordinate-based, and HGVS nomenclature-based inputs.

The major contribution of CNVision is not simply automation, but standardization. Intragenic CNV interpretation often requires repeated manual decisions about transcript selection, exon boundaries, coding regions, partial overlap, genome build, and reading-frame effect. CNVision formalizes these steps into a reproducible computational workflow, reducing the likelihood that different users will reach different conclusions from the same variant description.

### Comparison with existing workflows

Many existing genomic annotation tools are designed for broad variant interpretation, variant prioritization, pathogenicity prediction, or sequence-level annotation. These systems are valuable, but intragenic CNVs pose a narrower and technically distinct problem: determining how a deletion or duplication within a gene affects the coding sequence and reading frame. This requires accurate mapping between genomic coordinates, transcript structure, exon boundaries, coding sequence regions, and variant breakpoints.

Manual interpretation can resolve these questions, but it is time-consuming and inconsistent, highlighting the practical benefit of automated analysis demonstrated by CNVision (Figure 3). CNVision addresses this gap by focusing specifically on the molecular consequences of intragenic CNVs. Its input flexibility allows users to analyze variants described in the formats most commonly encountered in clinical reports, databases, and publications. Its output provides the computational basis for the predicted consequence rather than only a final label.

### Importance of base-pair-level interpretation

A central feature of CNVision is base-pair-level coding sequence overlap calculation. Exon-level descriptions are often insufficient because not all exonic sequences are coding, and not all CNVs affect complete exons. A CNV may overlap an untranslated region, part of a coding exon, several complete coding exons, or a combination of coding and non-coding sequence. Consequence prediction therefore requires more than counting affected exons.

By calculating the exact affected coding sequence length, CNVision can classify variants whose functional consequences depend on breakpoint precision. This is especially important for partial exon events, where a small difference in coordinate position may alter the predicted reading-frame effect. The framework’s deterministic calculation also makes the result easier to audit, since the affected coding length and modulo result can be directly reviewed.

### Discrepancies in published variant interpretation

The evaluation of published CNVs (Table 2) showed that CNVision identified a subset of variants whose predicted consequences differed from reported classifications. Manual re-evaluation supported the CNVision-derived classifications in these cases. This finding highlights a practical challenge in intragenic CNV interpretation: even when variants are reported in the literature, their functional consequences may be affected by transcript choice, coordinate interpretation, genome build conversion, or manual arithmetic error.

CNVision may therefore serve not only as an interpretation tool but also as a quality control framework. By applying the same rules consistently across CNVs, the system can identify variants that warrant reinspection. This is particularly useful when interpreting older reports, publications using different transcript references, or variant descriptions lacking complete breakpoint information.

### Strengths of the framework

CNVision has several strengths. First, it supports multiple input modalities, allowing users to analyze CNVs described by exons, genomic coordinates, or HGVS notation. Second, it uses transcript-aware coding sequence mapping rather than relying on gene-level or exon-count-based approximation. Third, it provides transparent intermediate calculations, including affected coding sequence length and reading-frame classification. Fourth, the application is web-accessible and containerized, making it easier to deploy and use across computational environments.

The framework is also intentionally focused. Rather than attempting to replace broader clinical interpretation systems, CNVision addresses a specific technical bottleneck in intragenic CNV consequence prediction. This focused scope makes the logic transparent and allows the tool to be integrated into larger research or clinical workflows as a functional consequence module.

### Limitations

Several limitations should be acknowledged. The current version of CNVision processes single-gene and single-variant inputs, which limits its utility for high-throughput genomic workflows. Many research and clinical pipelines generate large CNV callsets that require batch processing, filtering, and exportable results. Future versions should support CSV or VCF-like batch input formats to improve scalability.

The current interface also has a limited graphical representation of variant effects. Although CNVision reports exon coverage and coding sequence calculations, visual diagrams showing the gene model, affected exons, coding regions, and predicted consequences would improve interpretability. Such visualizations would be especially useful for users reviewing partial exon events or presenting variant interpretations in reports.

Transcript selection remains another important limitation. Although the use of standardized transcript resources improves consistency, some genes have multiple clinically relevant isoforms. A CNV may be in-frame in one transcript and frameshift in another, or may affect the coding sequence in one isoform but not another. Future versions should allow users to inspect alternative transcripts while still clearly identifying the primary reference transcript used for default classification.

Finally, CNVision predicts molecular consequences but does not independently determine clinical pathogenicity. A predicted frameshift may support loss-of-function interpretation, but clinical classification requires additional evidence, including disease mechanism, inheritance, variant frequency, segregation, phenotype match, and gene-disease validity. CNVision should therefore be used as a functional interpretation component within a broader variant classification workflow.

### Future development

Future development will focus on expanding CNVision from single-variant analysis to scalable CNV interpretation. Planned improvements include batch upload support, structured downloadable outputs, improved visualization of exon-level and coding sequence effects, and expanded support for complex variant structures. Integration with clinical and genomic databases may also allow CNVision outputs to be combined with gene-disease validity, ClinVar assertions, population data, and disease mechanism information.

## Conclusions

CNVision provides a reproducible, transcript-aware framework for functional interpretation of intragenic CNVs. By converting heterogeneous CNV descriptions into standardized coding sequence calculations, the framework enables consistent prediction of in-frame, frameshift, and non-coding consequences. Evaluation of 200 curated CNVs demonstrated complete concordance with manual transcript-level analysis across the tested dataset. CNVision addresses an important gap in current genomic workflows and provides a foundation for future development toward high-throughput, visually interpretable, and clinically integrated intragenic CNV analysis.

## Availability and Requirements

**Project name**: CNVision.

**Project home page**: https://github.com/mliao2710/CNVision

**Operating system**: Platform-independent through Docker; Linux deployment recommended.

**Programming language**: Python, HTML, CSS, Bootstrap.

**Frameworks and dependencies**: Flask, Docker.

**Input formats**: Exon-based CNV descriptions, genomic coordinate intervals/ISCN microarray nomenclature, and HGVS genomic notation.

**License**: MIT License

**Restrictions for use by non-academics**: No restrictions

**Web application**: https://cnvision-aqhgffh9etbchmc3.westus3-01.azurewebsites.net/

## Abbreviations

ISCN: International System for Human Cytogenomic Nomenclature
HGVS: Human Genome Variation Society
CNV: Copy Number Variant
CDS: Coding DNA Sequence
NMD: Nonsense-Mediated Decay
NGS: Next-Generation Sequencing
FISH: Fluorescence In Situ Hybridization
aCGH: Array Comparative Genomic Hybridization
WES: Whole-Exome Sequencing
WGS: Whole-Genome Sequencing
SNV: Single-Nucleotide Variant
MANE: Matched Annotation from NCBI and EMBL-EBI
RefSeq: Reference Sequence
ACMG: American College of Medical Genetics and Genomics
UCSC: University of California, Santa Cruz
HTML: HyperText Markup Language
CSS: Cascading Style Sheets
CSV: Comma-Separated Values

## Declarations

### Ethics approval and consent to participate

Not applicable. This study only used manually crafted and published variant examples and did not involve enrollment of human participants or generation of new patient-derived sequencing data.

### Consent for publication

Not applicable.

### Availability of data and materials

The CNVision source code is available at: https://github.com/mliao2710/CNVision. The curated validation dataset is available in the Supplementary Table. The software website: https://cnvision-24903.azurewebsites.net/.

### Authors’ contributions

M.L. completed all computer programming, website design, architecture development, validation testing, paper writing, and paper editing. X.Z. contributed to conceptualization, supervision, validation, data curation, coding guidance, and paper editing. All authors read and approved the final manuscript.

## Acknowledgements

Not applicable.

